# Genomic architecture and adaptive plasticity of *Enterococcus lactis* strains isolated from extreme Semi-arid environments

**DOI:** 10.64898/2026.06.05.730388

**Authors:** Javier Vanegas, C.M. Gaviria Prieto, Hillary Manotas

**Affiliations:** Faculty of Sciences, Universidad Antonio Nariño, Carrera 3 Este # 47A-15, Bogotá, Colombia; Colegio Mayor de Cundinamarca University, Bacteriology Program, Cl. 28, #5B-02, Bogotá, Colombia

**Keywords:** *Enterococcus*, agricultural biotechnology, abiotic stress, resource island, comparative genomics

## Abstract

The presence of *Enterococcus lactis* in semi-arid “resource islands” the remarkable ecological plasticity of a species often associated with host-related environments. Characterizing the genomic mechanisms that facilitate its persistence in extreme edaphic niches is crucial for exploring its biotechnological potential in arid agriculture. This study characterized the genomic architecture, abiotic stress tolerance, and plant growth-promoting (PGP) capabilities of six *E. lactis* strains isolated from the rhizosphere of *Pithecellobium dulce* and *Haematoxylum brasiletto* in La Guajira, Colombia. We compared the pangenomes of the isolates with clinical and environmental reference strains. Genomic predictions were validated through in vitro assays for thermal, saline, and pH stress, PGP traits, and biosafety (hemolysis, biofilm formation). Analysis revealed a pangenome with a conserved 2,113-gene core and a highly plastic 3,134-gene accessory genome. The core genome encodes robust machinery for osmotic stress (e.g., opuA–C operons) and DNA repair (uvrC), while the accessory genome is heavily shaped by Horizontal Gene Transfer, containing abundant Mobile Genetic Elements (6.3%–16.4%). Phenotypically, strains exhibited high resilience to heat (50°C), salinity (5% NaCl), and alkalinity (pH 12). Adaptation in these isolates favors metabolic parsimony: rather than complex phytohormone synthesis, the strains prioritize inorganic phosphate solubilization (conserved pst system) and harbor a complete 2,3-butanediol cluster for volatile-mediated plant interaction. Notably, strain IS_B39 produced siderophores and carried a specific RiPP-like biosynthetic cluster, indicating niche-specific functional diversification. Genomic and phenotypic screening confirmed a safe profile, lacking key virulence factors. These findings define a robust, low-risk genomic toolkit, supporting the potential of *E. lactis* as a tailored bioinoculant for sustainable agriculture in extreme, water-limited environments.

**Importance:** *Enterococcus* species are traditionally studied as clinical pathogens or dairy-associated bacteria, leaving their ecological role in natural, non-host environments largely overlooked. This study challenges conventional paradigms by exploring *Enterococcus lactis* strains naturally persisting in the extreme, water-limited soils of semi-arid “resource islands” in La Guajira, Colombia. Through functional genomics and laboratory validation, we demonstrated how these bacteria utilize a specialized genetic toolkit to withstand extreme heat and alkalinity, while actively promoting plant resilience. Rather than relying on complex hormone production, they optimize vital nutrient uptake like phosphorus. These findings significantly advance environmental microbiology by uncovering the hidden survival strategies of lactic acid bacteria in arid lands, showcasing their immense potential as sustainable bioinoculants to support global dryland agriculture under climate change stress.

## 1. Introduction

Semi-arid ecosystems impose severe selective pressures on microbial life, including chronic water limitation, high ultraviolet (UV) irradiance, and extreme thermal fluctuations (Pugnaire et al., 2004). In these heterogeneous landscapes, nurse plants facilitate the formation of “resource islands” (RIs) by modulating the microclimate, accumulating organic matter, and buffering soil temperature. These edaphic niches generate biogeochemical gradients that establish hotspots of biological activity and drive the recruitment of specific microbiomes (Pugnaire et al., 2004). Far from being passive residents, bacteria associated with RIs play a fundamental role in ecosystem functioning; their metabolic activities—such as nitrogen fixation and phosphate solubilization—are essential for maintaining soil fertility and supporting host plant resilience against abiotic stress (Armas et al., 2011; Sun et al., 2020). However, the assembly mechanisms of these microbiomes and the specific genomic traits that enable non-native taxa to colonize these competitive niches remain poorly characterized.

Among the diverse soil microbiota, the genus *Enterococcus* presents a compelling model for studying adaptation. While *Enterococcus lactis* has been historically characterized as a dairy-associated bacterium or a gastrointestinal commensal with high genomic plasticity (Morandi et al., 2013; Abril et al., 2022), its ecological competence in oligotrophic soil environments is paradoxical. Unlike typical rhizobacteria (e.g., *Bacillus* or *Pseudomonas*), enterococci generally lack robust biosynthetic pathways for amino acids and vitamins, relying instead on nutrient-rich host environments. Consequently, the isolation of *E. lactis* populations from the semi-arid RIs of the La Guajira Peninsula highlights their ecological versatility. While members of this species are frequently associated with gastrointestinal environments and fecal dispersal (Byappanahalli et al., 2012), their occurrence in these soils suggests a specialized metabolic repertoire that enables persistence under extreme edaphic conditions. Therefore, characterizing the genomic basis of their adaptation to free-living edaphic niches is crucial to understanding their ecological role in neotropical drylands.

This ecological resilience may be explained through the lens of microbial exaptation. Under this framework, traits originally selected for survival in diverse host-associated environments—such as osmotic stress tolerance—could be co-opted to confer fitness in saline or water-limited soils (Miller et al., 2020). However, beyond intrinsic resilience, successful colonization in these ecosystems likely requires functional specialization. As proposed by the functional integration hypothesis, edaphic colonizers may actively participate in soil biogeochemistry, potentially acquiring plant growth-promoting (PGP) traits via Horizontal Gene Transfer (HGT) to establish rhizosphere-associated niches (Bonatelli et al., 2021).

To explore the biotechnological potential and adaptive strategies of this species in extreme environments, this study characterizes the genomic and functional landscape of six *E. lactis* strains isolated from the rhizosphere of *Pithecellobium dulce* and *Haematoxylum brasiletto* in the semi-arid zone of La Guajira, Colombia. We employed a combined approach of Whole-Genome Sequencing (WGS), comparative pangenomics, and in vitro phenotypic validation. Our specific objectives were to: (i) define the adaptive genomic repertoire supporting abiotic stress tolerance and PGP activity; (ii) identify key metabolic genes that distinguish these environmental isolates; and (iii) assess their biosafety profile to evaluate their suitability for biotechnological applications..

## 2. Materials and Methods

### 2.1. Sampling and bacterial isolation

Sampling was conducted at the “Fundación Cerrejón para el Progreso de La Guajira” on the La Guajira Peninsula (approx. 11°32’N, 72°15’W), a semi-arid region in northern Colombia. The study targeted resource islands (Ris) associated with the nurse trees *Pithecellobium dulce* (site T) and *Haematoxylum brasiletto* (site B), including a bare soil control. Twelve composite soil samples were collected from a depth of <2 cm, covering the area from the trunk base to the canopy edge. The control sample was obtained 1.5 m from the RIs perimeter to represent unmodified bulk soil. One gram of each sample was enriched in Pseudomonas Agar F (PAF) medium (10 g L⁻¹ proteose peptone, 10 g L⁻¹ casein hydrolysate, 1.5 g L⁻¹ anhydrous MgSO₄, 1.5 g L⁻¹ K₂HPO₄, 10 mL L⁻¹ glycerol) to foster the recovery of diverse taxa (Aagot et al., 2001). Isolates displaying morphologies consistent with *Enterococcus* spp. were purified through three successive streaking rounds on Tryptic Soy Agar (TSA) at 30 °C (Ewald et al., 1992). Six isolates passing initial morphological and biochemical verification were selected for characterization and cryopreserved at −70 °C in 20% (v/v) glycerol.

### 2.2. DNA extraction, sequencing, and assembly

Genomic DNA was extracted from exponential-phase cultures using the DNeasy Power Soil Pro kit (QIAGEN, Germany) following the manufacturer’s protocol. DNA integrity and concentration were verified via 1% agarose gel electrophoresis and a Qubit fluorometer, respectively. Whole-Genome Sequencing (WGS) was performed on the Illumina NovaSeq 6000 platform, generating 2 × 150 bp paired-end reads with a minimum coverage of 100×. Raw reads were quality-checked using FastQC v0.11.9. Adapters and low-quality bases were removed using Trimmomatic v0.39 (Amuasi et al., 2023) with a sliding window Phred score filter of <30 and a minimum length of 50 bp. Genomes were assembled *de novo* using Unicycler v0.4.8. Assembly quality was evaluated using QUAST v5.2.0 (metrics: N50, contig count, genome size), and completeness/contamination were assessed using CheckM v1.2.4 based on lineage-specific marker genes (Parks et al., 2015).

### 2.3. Taxonomic identification and phylogenomic analysis

Taxonomic identification employed a polyphasic genomic approach. Species delineation was determined by Average Nucleotide Identity (ANI) using OrthoANIu (>95% threshold) (Yoon et al., 2017) and Digital DNA-DNA Hybridization (dDDH) using the Type Strain Genome Server (TYGS) (>70% threshold) (Meier-Kolthoff et al., 2022). Marker genes (*16S rRNA*, *gyrA*, *rho*) were validated against the NCBI database. The six isolates (IS_B17, IS_B38, IS_B39, IS_T25, IS_T26, IS_T34) were compared against two industrial *E. lactis* strains (15-3, MX2-1), the type strain *E. lactis* CX2-6_2, and the clinical reference *E. faecium* SRR24. Phylogenomic analysis was conducted on the BV-BRC v0.11.9 platform based on a MAFFT alignment of 1000 single-copy core genes (1,118,325 aligned nucleotides). Tree topology robustness was assessed using RAxML Fast Bootstrapping (N = 100).

### 2.4. Genome annotation and comparative pangenomics

Genomes were annotated using the BV-BRC Prokaryotic Genome Annotation Service. Functional assignments were refined by cross-referencing a custom database constructed through an exhaustive scientific literature search of experimentally validated genes associated with abiotic stress, plant growth-promoting (PGP) traits, and secretion systems. Comparison genomic analysis was performed using Roary v3.13.0 (Page et al., 2015) to delineate the core and accessory genomes of the La Guajira strains relative to reference genomes, focusing on the specific adaptive divergence of this edaphic clade. The distribution of stress resistance and interaction markers (Sec, Tat, T4SS) was visualized using TBtools (Chen et al., 2020). Biosynthetic Gene Clusters (BGCs) were identified using antiSMASH v8.0 (Blin et al., 2021), and Mobile Genetic Elements (MGEs) were predicted using IslandViewer 4 (Bertelli et al., 2017) to evaluate Horizontal Gene Transfer (HGT) potential.

### 2.5. In vitro phenotypic evaluation

All functional assays were performed in biological triplicate with uninoculated negative controls and reference positive controls (*Azospirillum brasilense*). Abiotic stress tolerance was evaluated by monitoring optical density (OD_600_) in LB broth over 48 h under modified conditions: temperature (50 °C and 70 °C), salinity (5%, 10%, and 15% NaCl), and pH (4.5, 9.0, 11.0, and 12.0). Regarding PGP traits, Indole-3-acetic acid (IAA) production was quantified in LB broth supplemented with 500 µg mL⁻¹ L-tryptophan using Salkowski’s reagent (Glickmann & Dessaux, 1995); siderophore production was assessed using the Chrome Azurol S (CAS) liquid assay (Schwyn & Neilands, 1987); phosphate solubilization was quantified in NBRIP medium via the molybdenum blue method (Nautiyal, 1999); and nitrogen fixation potential was evaluated qualitatively in Nfb medium using bromothymol blue as a pH indicator (Buisset et al., 2025). Finally, the biosafety profile was determined *in silico* using PathogenFinder 1.1 and validated phenotypically by testing hemolytic activity on 5% sheep blood agar, erythrocyte agglutination, and biofilm formation in 96-well plates using crystal violet staining (OD_590_) (Tula et al., 2022). Quantitative data were expressed as the mean ± standard deviation (SD). For stress tolerance assays, growth was scored as positive (+) or negative (−) based on the negative control.

## 3. Results

### 3.1. Isolation and phylogenomic identification

The six isolates (IS_B17, IS_B38, IS_B39, IS_T25, IS_T26, and IS_T34) were characterized as Gram-positive cocci with circular, off-white colony morphology. Phylogenomic analysis based on 1000 single-copy core genes positioned all isolates within a distinct, highly supported monophyletic clade (Bootstrap = 100) nested within the *Enterococcus lactis* species group (Fig. 1A). This clade was clearly separated from the clinical reference *E. faecium* SRR24 and outgroup genera. Taxonomic assignment was confirmed by genomic identity metrics (Fig. 1B); all isolates exhibited Average Nucleotide Identity (ANI) values between 97.8% and 98.6%, and Digital DNA-DNA Hybridization (dDDH) probabilities >93% relative to the *E. lactis* type strain (CX 2-6_2). These values exceeded the established species delineation thresholds of 95% for ANI and 70% for dDDH.

**Figure 1.**
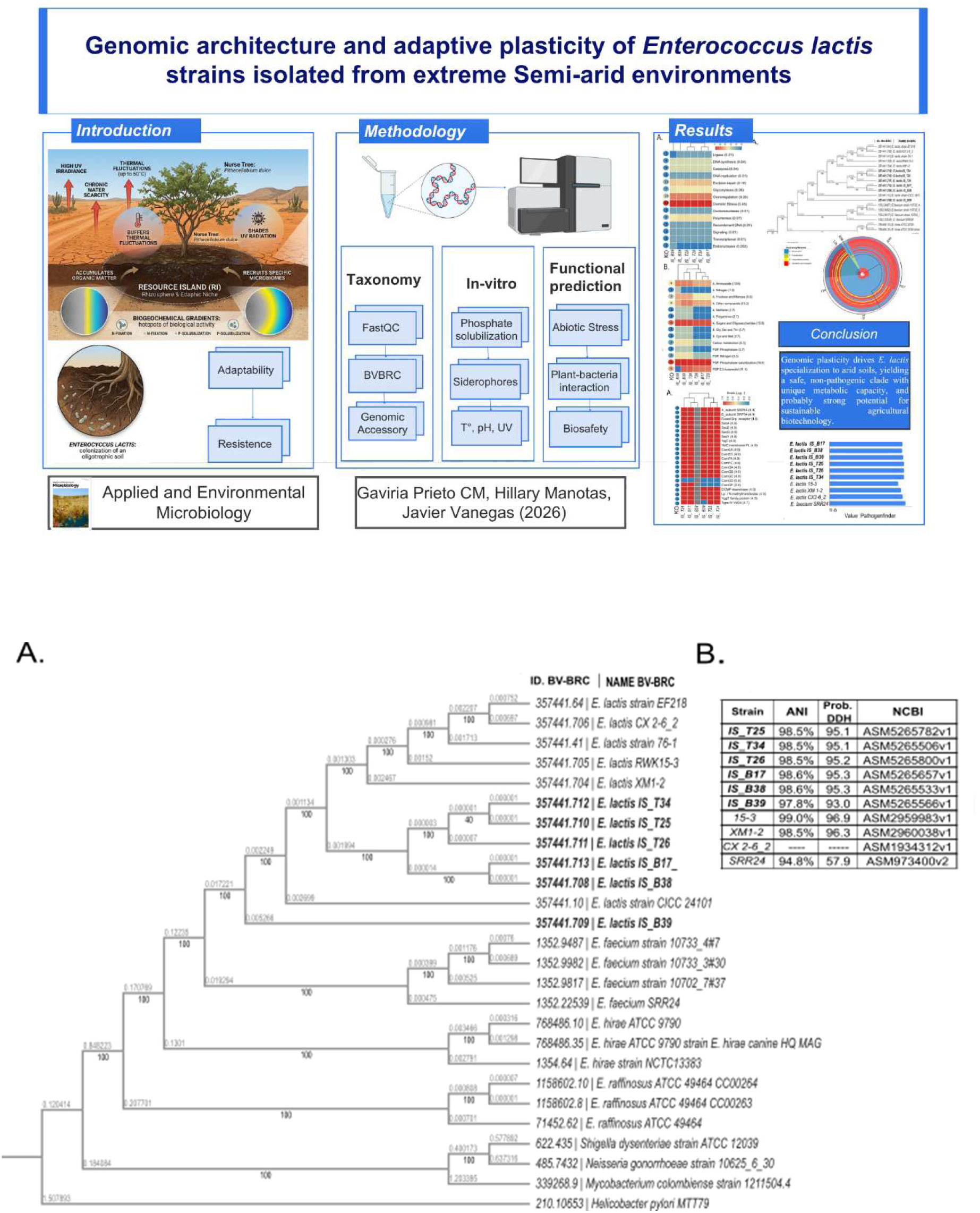
Taxonomic delimitation of the *Enterococcus lactis* strains from La Guajira. (A) Maximum Likelihood phylogenetic tree based on the alignment of 2,113 core-genome genes. Bootstrap support values (>50%) from 1,000 replicates are shown at the nodes. Strains isolated in this study are prefixed “IS_”. Reference *E. lactis* genomes from diverse origins and an *Enterococcus faecium* outgroup were included. The scale bar represents nucleotide substitutions per site. (B) Table of Average Nucleotide Identity (ANI) and digital DNA-DNA Hybridization (dDDH) values calculated against the *E. lactis CX2-6_2* reference genome. The results confirm that the La Guajira strains form a monophyletic clade and meet the genomic criteria for classification as *E. lactis*.

### 3.2. Genome assembly and features

*De novo* assembly yielded high-quality draft genomes ranging from 2.73 to 2.81 Mbp (Table 1). The assemblies displayed low fragmentation (18 to 57 contigs) and N50 values between 235 and 274 kbp. Genome completeness was estimated at >99% with minimal contamination (<0.3%) for all strains. The G+C content was conserved (38.0–38.3%). Annotation predicted between 2,632 and 2,840 protein-coding sequences (CDS) per genome, with approximately 28% classified as hypothetical proteins. Each genome contained 44 to 46 tRNA genes and 3 complete rRNA operons.

**Table 1.** Assembly statistics and genomic features of the six E. lactis strains.

| Characteristic | *IS_B17* | *IS_B38* | *IS_B39* | *IS_T25* | *IS_T26* | *IS_T34* | *CX 2-6_2* |
| --- | --- | --- | --- | --- | --- | --- | --- |
| **Assembly Quality** |  |  |  |  |  |  |  |
| Contigs (No.) | 38 | 41 | 57 | 18 | 24 | 18 | 1 |
| N50 (kbp) | 264.6 | 235.3 | 269.6 | 274.8 | 274.8 | 274.8 | 2559.2 |
| L50 (No.) | 4 | 4 | 4 | 4 | 4 | 4 | 1 |
| Completeness (%) | 99.42 | 99.42 | 99.12 | 99.42 | 99.42 | 99.42 | 99.32 |
| Contamination (%) | 0.27 | 0.27 | 0.21 | 0.27 | 0.27 | 0.27 | 0.27 |
| Genome fraction (%) | 85 | 85 | 85 | 84 | 84 | 84 | - |
| Indels per 100 kbp | 39 | 39 | 55 | 35 | 38 | 38 | - |
| **Genomic Features** |  |  |  |  |  |  |  |
| Genome size (kbp) | 2813 | 2812 | 2791 | 2725 | 2752 | 2725 | 2559 |
| G+C content (%) | 38.1 | 38.1 | 38.2 | 38.3 | 38.3 | 38.3 | 38.27 |
| Total CDS (No.) | 2724 | 2728 | 2840 | 2632 | 2669 | 2635 | 2505 |
| tRNA (No.) | 44 | 44 | 46 | 46 | 46 | 46 | 67 |
| rRNA (No.) | 3 | 3 | 3 | 3 | 3 | 3 | 18 |
| **Functional Novelty** |  |  |  |  |  |  |  |
| Hypothetical CDS (No.) | 777 | 782 | 792 | 705 | 737 | 713 | 614 |
| Hypothetical CDS (%) | 28.5 | 28.7 | 27.9 | 26.8 | 27.6 | 27.1 | 26.3 |

The table summarizes key metrics for assembly quality and genome characteristics for the isolates from La Guajira and the reference strain *E. lactis* CX 2-6\_2 (NCBI accession: NZ\_CP079880.1). Quality metrics estimated using CheckM.

### 3.3. Comparative genomics

Genomic analysis of the six La Guajira strains revealed a core genome of 2.113 genes, representing conserved metabolic and cellular functions. The accessory genome comprised 3.134 genes, constituting the primary source of genetic diversity within the clade (Fig. 2). Functional categorization (Table 2) showed that while core metabolic genes were evenly distributed, the accessory genome was enriched in genes related to environmental interaction, specifically secretion systems and transport mechanisms (Table S1).

**Figure 2.**
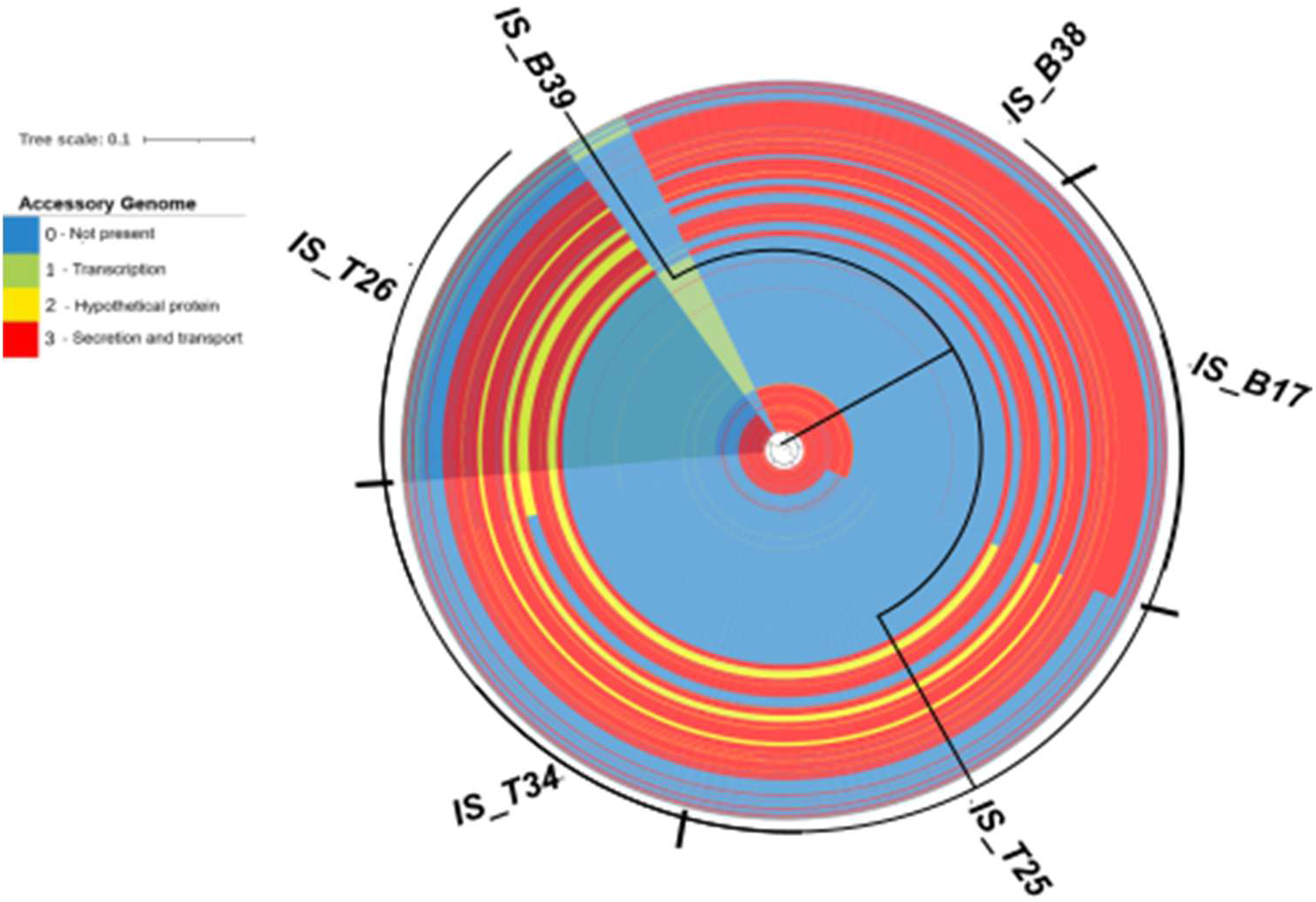
Comparison genome visualization of the six *E. lactis* strains from La Guajira. A core-genome phylogenetic tree is shown at the center. Each concentric ring represents a strain’s genome, with colored segments indicating the presence of gene clusters from the 3,134-gene accessory genome. The color legend denotes functional categories: Red (Secretion/Transport), Yellow (Hypothetical Proteins), Green (Transcription). The figure highlights the high genomic plasticity driven by the accessory genome, with a notable enrichment of genes involved in environmental interaction.

**Table 2.**
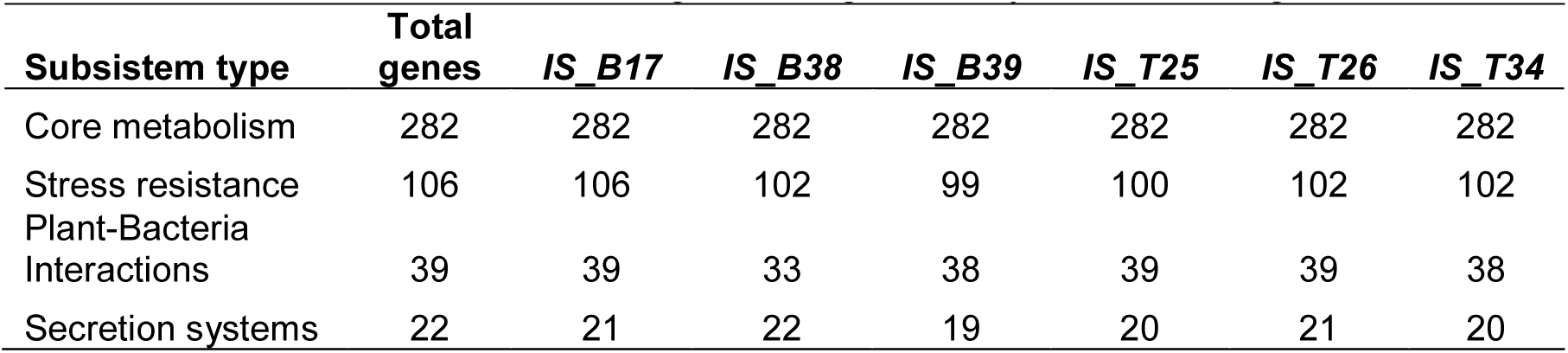
The table shows the number of genes assigned to key functional categories.

### 3.4. Genomic potential for stress adaptation and plant interaction

Functional annotation identified a conserved genetic repertoire for abiotic stress resistance (Fig. 3A, Table S2). All strains harbored the complete osmoregulation machinery, including the *opuA–C* and *proA–C* operons for osmoprotectant uptake. Regarding DNA repair, strains *IS_B17* and *IS_B38* were distinct from the other isolates, carrying additional copies of key repair genes such as *uvrC* and *mutS* (Fig. 3A).

**Figure 3.**
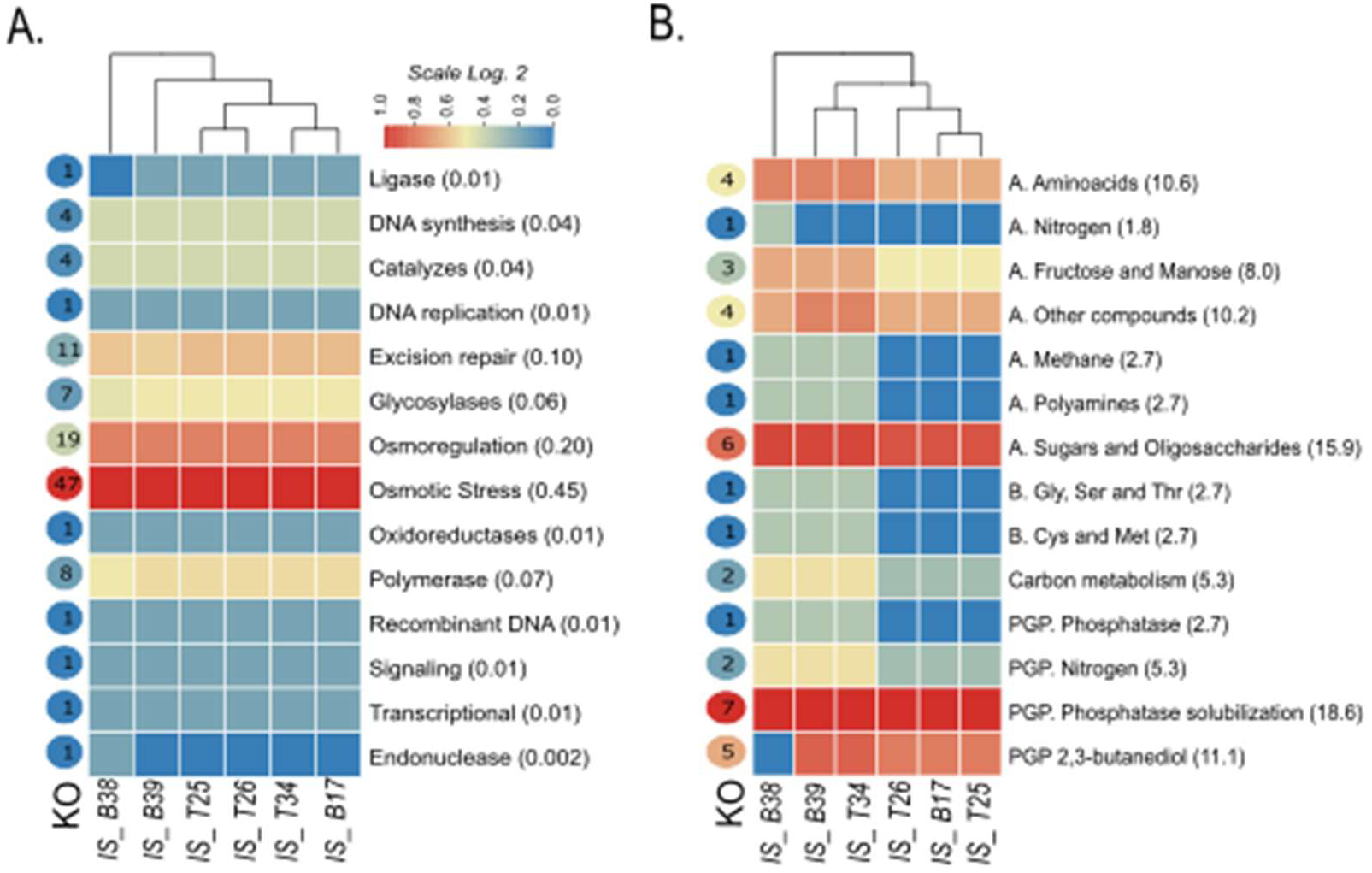
Heatmap analysis of genes related to abiotic stress and plant growth promotion. The heatmaps visualize the relative abundance (log₂ normalized) of genes in each functional category. Red indicates above-average abundance, and blue indicates below-average abundance. (A) Abiotic Stress Resistance Profile. All strains share a strong genetic signature for “Osmotic Stress” and “Osmoregulation.” *E. lactis* B17 and *E. lactis* B38 cluster together due to a larger repertoire of DNA repair genes. (B) Plant Growth Promotion (PGP).

In terms of plant-bacterium interactions, the core genome of all strains encoded the high-affinity phosphate transport system (*pstA/B/C/S*) and enzymes for the catabolism of plant-derived sugars (Fig. 3B, Table S3). Notably, a complete biosynthetic gene cluster (BGC) for the production of the volatile compound 2,3-butanediol (*alsS*, *alsD*, *ilvB*) was consistently identified in all six genomes (Table S4). Analysis of Mobile Genetic Elements (MGEs) revealed a prevalence of insertion sequences and prophages, ranging from 6.3% to 16.4% of the total genome size. Strain *IS_B39* exhibited the highest MGE content (16.4%) and harbored a unique genomic island containing a RiPP-like (Ribosomally Synthesized and Post-translationally Modified Peptide) BGC, a trait absent in the other isolates (Table S5).

### 3.5. Phenotypic validation

*In vitro* assays confirmed the genomic predictions regarding stress tolerance (Table 3). All strains demonstrated growth at 50°C and in the presence of 5% NaCl. Furthermore, strains IS_B38 and IS_T25 supported growth at 70°C. Strain *IS_T34* exhibited tolerance to alkaline conditions (up to pH 12). However, growth was inhibited under extreme stress conditions (10% NaCl, or pH 3). Regarding PGP traits, no nitrogenase activity or IAA production was detected in any strain; however, the positive control strain *A. brasilense* did show a positive result for both activities. Siderophore production was exclusively observed in strain *IS_B39*, correlating with its unique genomic features. Phosphate solubilization was positive for four strains.

**Table 3.**
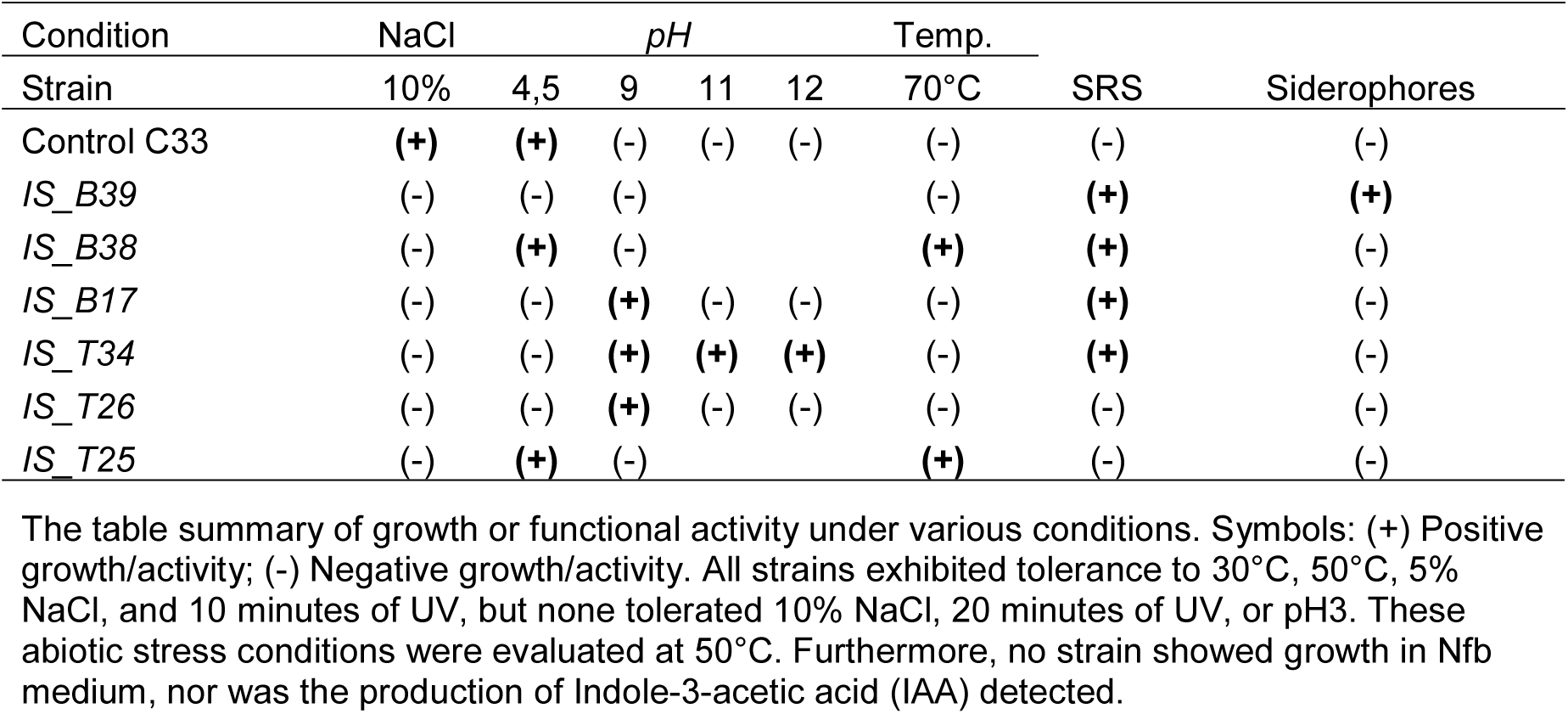
Phenotypic profile of *E. lactis* strains under abiotic stress and for PGP activity.

### 3.6. Biosafety profile

*In silico* analysis using PathogenFinder predicted a low pathogenic potential for all isolates (score <0.7), grouping them with non-clinical food-grade strains and distinct from the clinical reference *E. faecium* SRR24 (Fig. 4B). Genomic screening confirmed the absence of key virulence factors (e.g., *gelE*, *esp*) and antibiotic resistance markers (Table S6). Phenotypically, all six strains were non-hemolytic (gamma hemolysis) on sheep blood agar, coagulase-negative, and unable to form biofilms under the tested conditions.

**Figure 4.**
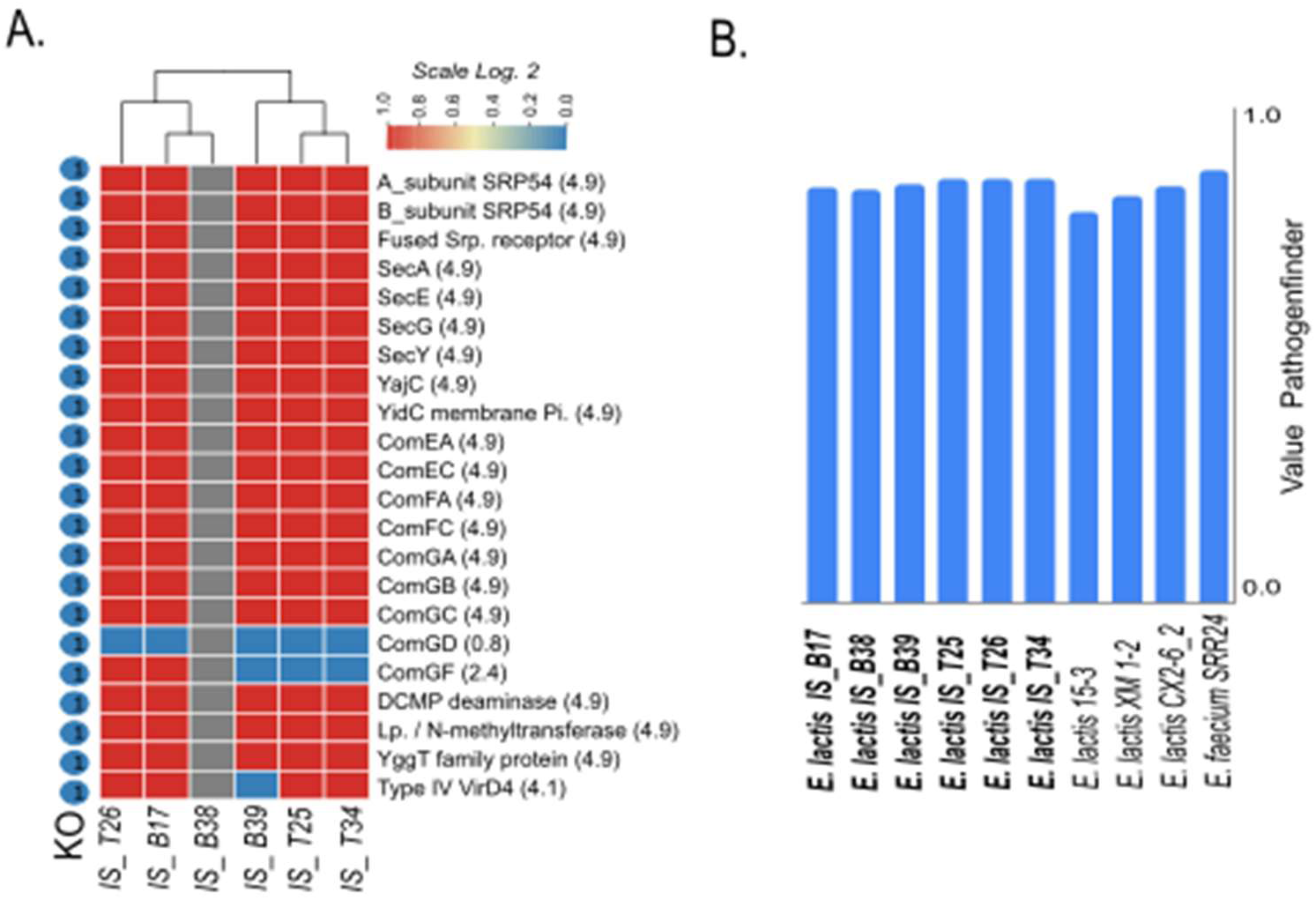
Integrated analysis of secretion repertoire and genomic biosafety profile. (A) Secretion and transport profile. The heatmap shows the abundance of key genes in secretion and transport systems, revealing a highly conserved potential for environmental interaction via the general secretion (SEC, SRP) and natural competence (Type II) pathways. (B) Genomic Biosafety Profile. The bar graph displays the pathogenicity score predicted by PathogenFinder. The La Guajira strains (score < 0.7) cluster with non-clinical isolates, showing a significantly lower risk profile than the clinical reference *E. faecium SRR24* (score > 0.7). Detailed gene information is in Supplementary File S6.

## 4. Discussion

### 4.1 Genomic identification of *E. lactis* strains

The confirmed presence of *Enterococcus lactis* in the RIs of La Guajira (Fig. 1) challenges the species’ established ecological paradigm. The genomic architecture of these isolates suggests a dual functional strategy that facilitates their persistence in extreme edaphic niches (CheckM analysis >99%, Table 1). This framework is based on an intrinsic stress-tolerance toolkit within the core genome, which provides the essential genetic repertoire for primary survival in the soil matrix. The core genome, evolved under the selective pressures of the intestine, conferred a robust machinery for osmotic tolerance and DNA repair (Gaca & Lemos, 2019), which were critical prerequisites for overcoming the stringent environmental filter of the soil. Upon this pre-existing platform of resilience, a second phase of edaphic specialization occurred, driven by the plasticity of the accessory genome. This adaptation to the RIs niche is evident in the acquisition of plant-growth-promoting (PGP) traits via HGT and in the substantial repertoire of hypothetical proteins (28%), which likely encode undescribed functional innovations (Cotta et al., 2025). Therefore, this model not only organizes our findings but also demonstrates the ecological plasticity of *E. lactis*, highlighting its ability to function as a robust edaphic competitor in extreme environments.

Notably, the monophyletic clustering of the La Guajira strains (Fig. 1) supports this model, suggesting that their edaphic specialization is the result of *in situ* diversification from a common ancestor that successfully navigated the initial transition (García-Solache & Rice, 2019).

### 4.2. Core genome as a prerequisite for survival

The core genome of the La Guajira isolates constitutes a robust functional foundation for abiotic stress resilience. Our analysis identified a core of 2,113 genes, indicating strong functional conservation. This substantial figure, when compared to the 1616 gene core identified in a more diverse set of 6 *E. lactis* strains (Lu et al., 2023), suggests strong functional conservation within our edaphic clade. This conserved genetic repertoire functions as a pre-adapted toolkit, originally selected not for the soil but by the selective pressures of the gastrointestinal niche (García-Solache & Rice, 2019).

This intrinsic resilience manifests in three critical functional areas. First, adhesion, a critical trait for persistence in the intestinal tract, is encoded in the core genome, as demonstrated by the presence of adhesion factors like *efaA* (Eaton & Gasson, 2001), a gene also reported as conserved in other *E. lactis* strains (Lu et al., 2023). Second, osmotic and chemical stress tolerance is a key feature. The machinery for osmoprotectant uptake (*opuA-C*, *proA-C*), essential for withstanding high concentrations of bile salts, was directly co-opted for survival in a soil with high salinity and hydric fluctuations. Third, the conserved repertoire for DNA repair (*UvrC, MutS*) (Fig. 3A) is a fundamental pre-adaptation for mitigating genotoxic damage, a function as relevant against oxidative stress from the host immune system as it is against the high UV irradiance of the desert (Gaca & Lemos, 2019). Therefore, this conserved suite of traits—adhesion, resilience, and repair—does not represent a *de novo* adaptation to the soil. It constitutes a fundamental physiological platform; these conserved traits—adhesion, resilience, and repair—provide the baseline robustness required to overcome the stringent selective filters of semi-arid soils. (Byappanahalli et al., 2012).

### 4.3. Edaphic specialization

The functional diversification of *E. lactis* is driven by its accessory genome (3,134 genes), which serves as a modular and adaptive genetic reservoir for biotechnological innovation. This locus of diversification is shaped by selective pressures on two interconnected fronts: abiotic resilience and biotic competitiveness. The primary selective filter in La Guajira is abiotic pressure. Our data demonstrate how the accessory genome provides direct genetic solutions, such as the gene-dosage effect observed in the overrepresentation of DNA repair genes (*UvrC, MutS*) in strains IS_B17 and IS_B38 (Fig. 3A, Table 3). This capacity is a fundamental mechanism for maintaining genomic integrity against the high mutational load imposed by UV irradiance and desiccation stress (Choudhary et al., 2016).

However, abiotic resilience is insufficient for ecological persistence. The second function of this genomic repertoire is to equip *E. lactis* for biotic interaction. Here, the genomic evidence points toward a complex strategy of communication and chemical antagonism. The conservation of BGCs for the production of cyclo-lactone-type autoinducers across the entire clade (antiSMASH, Table S5) suggests a basal capacity for intercellular communication (quorum sensing). Building on this foundation, specialization emerges: strain *IS_B39* harbors a unique RiPP-like BGC (Table S4), indicating the potential to produce antimicrobial peptides as a mechanism of competitive exclusion by interference, which is crucial in the competition for the limited resources of the RIs (Arias-Orozco et al., 2024).

The presence of these specialized modules, particularly the unique RiPP-like BGC in strain IS_B39, highlights the bioprospecting potential of these isolates for developing novel bio-based agricultural inputs. The evidence strongly points to Horizontal Gene Transfer (HGT), the high frequency of which is inferred from the notable abundance of Mobile Genetic Elements (MGEs) (6.3% to 16.44%) and the consistent detection of Genomic Islands (GIs) (IslandViewer4, Table S4) (Cotta et al., 2025). The IS_B39 strain is a paradigmatic example of this process; its genome, which contains the highest percentage of MGEs in the clade (16.44%), also harbors the most specialized functional repertoire, including the exclusive production of siderophores and a unique RiPP-like BGC. Therefore, strain IS_B39 is not merely a variant but exemplifies the process by which high genomic plasticity directly translates into tangible phenotypic innovation. HGT enables the incorporation of functionally cohesive and evolutionarily tested loci, significantly accelerating the rate of adaptation compared to vertical evolution (Arnold et al., 2022). Moreover, the bacterium possesses the machinery to actively facilitate this gene flow. The presence of the natural competence system (*comEA*, *comEC*) allows for the internalization of environmental DNA, while the components of a Type IV Secretion System (*virD4*) suggest a potential for conjugation. Collectively, this high genomic plasticity positions *E. lactis* as a highly adaptable ecological competitor, capable of actively remodeling its genome to optimize its fitness against the multiple challenges of an extreme environment (Wallden et al., 2010).

The long-term persistence of *E. lactis* in the RIs of La Guajira is underpinned by a strategic duality that balances energy conservation with habitat modification. The first pillar is a pronounced metabolic parsimony, an adaptive strategy where fitness is maximized through resource-use efficiency rather than growth rate (Allison, 2014). The genomic evidence for this strategy is compelling. The genome harbors an extensive repertoire for the catabolism of complex organic substrates, including plant-derived carbohydrates (*araA, galE, lacZ, xylA*) and amino acids (*serA, glyA, glnA*) (Fig. 3B). This catabolic versatility minimizes reliance on costly *de novo* biosynthesis. Critically, this efficiency strategy extends to phosphorus acquisition. The genome encodes the complete machinery of the high-affinity phosphate transport system (*PST; pstA/B/C/S, phoR/H/U*) (Fig. 3B), a mechanism that enables the active uptake of inorganic phosphate, an energetically more favorable alternative than the mineralization of organic phosphate (Santos-Beneit et al., 2008).

The phenotypic validation reveals more than a simple genotype-phenotype correlation, uncovering a layer of functional complexity and specialization within the La Guajira clade. The observed heterogeneity, such as the exclusive siderophore production by strain *IS_B39* (Table 2), despite the genomic potential for plant-bacterium interaction being comparable across other strains (Fig. 3B), suggests the existence of differential regulatory networks that orchestrate the expression of these traits. This pattern of functional specialization indicates that the clade, far from being a homogeneous population, has diversified its strategies to exploit specific micro-niches (Maestre et al., 2024). For instance, the superiority of strains *IS_B17* and *IS_T34* under alkaline conditions (Table 2) could reflect an adaptation to soil zones with high mineralization, while the thermotolerance of *IS_B38* and *IS_B39* might confer an advantage in the more superficial and exposed soil layers. This phenotypic divergence is the manifestation of the edaphic adaptation phase of these microorganisms (Rajakaruna 2018).

Even more revealing, from an eco-evolutionary perspective, is the universal absence of certain canonical PGP traits. Neither the production of IAA nor the capacity for nitrogen fixation was detected (Table 2). We interpret this absence not merely as a lack of genes, but as a reflection of phylogenetic constraints that favor alternative resource mobilization strategies. The fixation of N₂ gas is one of the most energetically costly biological processes (Sheoran et al., 2021), and Enterococcus lineages generally lack the *nif* machinery required for this function. Consequently, natural selection in oligotrophic environments likely favored strains that optimized low-cost resource mobilization, prioritizing the solubilization of inorganic phosphate—a universal trait in the clade and abundantly represented in its genome (Fig. 3B)—and iron chelation. This strategy, grounded in metabolic efficiency, represents an efficient adaptive solution, well-suited to the challenges of an environment characterized by extreme resource scarcity (Zhu et al., 2024).

The energy conserved through this metabolic parsimony is selectively invested in a second strategic pillar: niche construction (Laland et al., 2019). The complete biosynthetic pathway for 2,3-butanediol (*alsS, alsD, BudA, BudB, ilvB*) (Fig. 4) is the central mechanism of this strategy. This Volatile Organic Compound (VOC) has the potential to induce systemic stress tolerance in the host plant (Ryu et al., 2003; Mai et al., 2021). By possessing the genetic capacity to emit it, *E. lactis* could reinforce the resilience of the nurse tree upon which it depends. This establishes a potential positive feedback loop between the plant and the microbiome, where increased host stability translates into greater stability and predictability of the microbial habitat (Bever et al., 2012). Therefore, this equilibrium between strict energy conservation (parsimony) and selective investment in habitat stabilization (niche construction) constitutes the organizing principle of its resource allocation strategy. His metabolic parsimony, combined with high-affinity phosphate uptake (PST system), suggests that these strains are optimized for nutrient-poor soils, making them ideal candidates for sustainable fertilization in degraded lands.

### 4.4. Biosafety Profile

The favorable biosafety profile of the La Guajira strains is not an incidental finding but the predictable consequence of their niche-specific genomic profile, a finding that aligns with growing evidence distinguishing *E. lactis* from its pathogenic relatives. The genomic evaluation places this edaphic clade at the lower end of the risk spectrum (PathogenFinder score < 0.7) (Fig. 4B) (Mashzhan et al., 2025). This favorable profile, characterized by the absence of canonical virulence factors and biofilm-forming capacity, supports the safety of these strains for large-scale agricultural release. This genomic profile is notably consistent with recent safety evaluations of *E. lactis* strains from diverse origins, which also report a lack of this virulence arsenal (Lu et al., 2023).

This genomic prediction is strongly corroborated at the functional level. Phenotypic assays confirmed that our strains are universally non-hemolytic (gamma hemolysis) and coagulase negative, a safety phenotype that aligns with that of other *E. lactis* strains considered safe (Lad et al., 2022; Argemi et al., 2019). A critical and revealing point of divergence, however, is the inability of our strains to form biofilms. While the ability to form biofilms has been observed in other *E. lactis* strains (Lu et al., 2023), its absence in the La Guajira clade supports a favorable biosafety profile, as biofilm formation is a key virulence factor in clinical settings. Ecologically, this trait suggests a lifestyle trade-off: while lacking biofilm might reduce desiccation protection, it may favor dispersal over sessile persistence in this specific dryland context (Zhu et al., 2024).

Collectively, this integrated profile reinforces the argument that risk assessment in *Enterococcus* must be both species and clade specific. The genomic characterization of *E. lactis* strains from La Guajira RIs resource islands identified diverse abiotic stress resistance mechanisms in the core genome and plant growth-promoting traits enriched in the accessory genome (Lu et al., 2023; Maguvu et al., 2021). This dual genomic architecture supports both the intrinsic resilience hypothesis through conserved stress tolerance genes enabling soil colonization and the functional integration hypothesis through acquired PGP capabilities facilitating active participation in RIs biogeochemistry (Muller et al., 2001). These findings establish the biotechnological potential of strains from RIs as multifunctional bioinoculants for sustainable agriculture in semi-arid environments (Morandi et al., 2013; Choudhary et al., 2016).

### 4.5. Limitations and future perspective

The conclusions drawn from this study, while robustly supported by integrated genomic and phenotypic data, necessarily generate a new set of hypotheses. First, we acknowledge that our genomic model is derived from a geographically defined set of isolates from La Guajira; broader sampling efforts across diverse arid ecosystems will be valuable to confirm the universality of this adaptive trajectory. Additionally, while this work establishes a high-resolution map of genetic potential, it does not capture the dynamics of gene expression *in situ*. Validating our adaptive model will be essential through transcriptomic (RNA-seq) and metabolomic analyses (Cao et al., 2022). Such studies are crucial to capture the regulatory networks governing the activity of these strains, particularly in response to the specific biogeochemical cues provided by the nurse tree (Maestre et al., 2024).

Furthermore, our *in vitro* assays represent a necessary simplification of the complex ecological interactions within the RIs soil. The translation of the observed traits into ecologically significant benefits must be rigorously evaluated in controlled mesocosms and field settings (Albornoz et al., 2022). Finally, the substantial fraction of unannotated genes (∼28%) represents a current analytical limitation but also a significant scientific opportunity. This unexplored functional landscape likely harbors novel proteins underpinning the unique adaptation of the La Guajira clade (Vincent, 2024). Elucidating the function of these genes through targeted functional genomics will be fundamental to uncovering novel biotechnological targets.

## 5. Conclusion

This research successfully characterized *E. lactis* strains isolated from La Guajira, providing a robust genomic framework that explains their functional adaptation to semi-arid ecosystems. Our findings establish that their survival in these extreme environments is driven by a dual strategy: a conserved core genome that ensures high resilience to abiotic stress (validated by tolerance to 50 °C and 5% NaCl), and a highly plastic accessory genome that enables specialized resource acquisition.

Functional analysis reveals a metabolic prioritization in these isolates, where habitat engineering via volatile organic compounds (specifically the 2,3-butanediol pathway) and low-cost nutrient mobilization are favored over energy-intensive synthesis pathways. This efficiency is a key trait for persistence in nutrient-limited edaphic niches.

From a biotechnological perspective, these findings demonstrate that *E. lactis* strains from extreme environments possess a unique ‘genomic toolkit’ of resilience and efficiency. Coupled with a confirmed biosafety profile (absence of virulence factors and biofilm formation), these isolates emerge as high-potential, low-risk candidates for the development of tailored bioinoculants, offering a sustainable solution for agriculture in water-scarce regions.

## Acknowledgements

We extend our sincere gratitude to Nelson Valero, Daiver Pinto, and Juan Taborda for their invaluable technical assistance and field support. We are profoundly grateful to the staff of the Cerrejón Foundation for the Progress of La Guajira (*Fundación Cerrejón Para El Progreso De La Guajira*) for facilitating access to the farm and supporting the sampling activities. We also thank Professor Yuly Bernal from the basic sciences laboratory for her support in providing supplementary materials.

## Appendix A. upplementary material

The following are the Supplementary data to this article.

## Data availability

The Whole-Genome Sequencing (WGS) data generated and analysed in this study have been deposited under BioProject <u>1127038</u> at the National Center for Biotechnology Information (NCBI). All assembled genomes, raw sequence reads, and associated metadata are publicly available via the links provided in the BioProject.

